# Magnetic Particle Imaging is a sensitive *in vivo* imaging modality for the quantification of dendritic cell migration

**DOI:** 10.1101/2021.09.22.461401

**Authors:** Julia J. Gevaert, Corby Fink, Jimmy Dikeakos, Gregory A. Dekaban, Paula J. Foster

## Abstract

Immunotherapies, such as dendritic cell- (DC-)based therapies, are useful for treating cancer as an alternative to or in combination with traditional therapies. Cells must migrate to lymphoid organs to be effective and the magnitude of the ensuing T cell response is proportional to the number of lymph node-migrated DC. With less than 10% of cells expected to reach their destination, there is a need for an imaging modality capable of sensitively and quantitatively detecting cells. MRI has been used to track DC using iron and 19F methods, with limitations. Quantification of iron-induced signal loss is indirect and challenging; 19F signal is directly quantifiable but lacks sensitivity. Magnetic Particle Imaging (MPI) directly detects superparamagnetic iron oxide nanoparticles (SPIO) and enables quantitation of low numbers of SPIO-labeled cells. Here we describe the first study using MPI to track and quantify the migration of DC, injected into the footpads of C57BL/6 mice, to the popliteal lymph nodes (pLNs). As DC migrate from the site of injection to the lymph nodes, we measured a decrease in signal in the footpads and an increase in signal at the pLNs. The presence of SPIO-labeled DC in nodes was validated by *ex vivo* MPI and histology. By measuring the iron mass per cell in samples of labeled cells, we were able to provide an estimate of cell number for each source of signal and we report a sensitivity of approximately 4000 cells *in vivo* and 2000 cells *ex vivo*. For some mice, MPI was compared to cellular MRI. We also bring attention to the issue of resolving unequal signals within close proximity, a challenge for many pre-clinical studies using a highly concentrated tracer bolus that over shadows nearby lower signals. This study demonstrates the clear advantage of MPI to detect and quantify cells *in vivo*, bridging the gap left by cellular MRI, and all other *in vivo* imaging modalities, and opening the door for quantitative imaging of cellular immunotherapies.

## Introduction

Cell therapy, an umbrella term encompassing the therapeutic administration of immune cells (such as T cells or dendritic cells (DC)) and stem cells (such as mesenchymal stem cells or pluripotent stem cells), is now recognized as a safe and effective approach to the treatment of a number of cancers^1^ as well as a growing list of autoimmune,^2^ degenerative,^3^ genetic^4^ and infectious diseases.^5^

Cancer immunotherapy is an innovative and evolving research area that involves the use of one’s own immune system to combat cancer. One approach is to target tumors with a tumor antigen-specific vaccine.^6,7^ A refinement of this approach uses antigen presenting cells, such as DC, which are particularly adept at triggering primary immune responses, as an adjuvant system for tumor antigen presentation. Over 300 DC cancer vaccine trials were conducted over the last 20 years and some products have received special regulatory approval frameworks by the U.S. Food and Drug Administration (FDA) and European Medicines Agency.^8^

Despite its immense potential, clinical results from DC cancer vaccine trials have been variable and discordant because of disparities in cell source, preparation, route of administration and implantation methodologies. It remains difficult to spatially and quantitatively track the migration of cells and the immune response that follows. Many critical questions about the delivery, anatomical location/distribution, numbers and persistence of DC remain unanswered. For example, in DC cancer immunotherapy, the magnitude of an anti-tumor response is proportional to the quantity of antigen presenting cells that reach a target lymph node,^9–11^ and therefore, it is crucial to know whether the injected cells have migrated to the target and in what magnitude to establish correlates with vaccine efficacy.

Magnetic resonance imaging (MRI) has been used to track DC migration. This involves labeling DC with superparamagnetic iron oxide nanoparticles (SPIO) prior to their administration and imaging with pulse sequences sensitive to iron. In MRI, SPIO produce negative contrast or signal loss, that is indirectly detected through its relaxation effects on protons. SPIO-based MRI cell tracking has very high sensitivity (10’s - 100’s of cells), albeit with low specificity, making it very difficult to accurately quantify the local tissue concentration of SPIO particles.^12^ Magnetic Particle Imaging (MPI) is a novel tracer-based imaging modality that directly detects the response of SPIO to an applied magnetic field.^13,14^ MPI has potential for overcoming the challenges of MRI-based cell tracking because it has high specificity (direct detection of only the response of SPIOs) and high sensitivity to nanogram quantities of SPIOs, which translates to hundreds of cells.^15^ Importantly, the MPI signal is linearly related to iron mass allowing for quantitation of the amount of iron from images, and with knowledge of the amount of iron/cell after SPIO labeling, the cell number can be calculated.^16^ The main limitation of MPI is resolution. Sensitivity and resolution are both determined in part by the type of SPIO employed for cell labeling. A second limitation is the ability to quantify two sources of MPI signal in close proximity, due to highly concentrated signal overshadowing lower tracer concentrations located nearby. This limits the dynamic range of the system when several samples with different concentrations are being imaged.^17^

In this study we evaluate the use of MPI for quantitative tracking of DC in a preclinical mouse model. In this model system, DC are administered subcutaneously into the mouse footpad and migrate to the draining lymph node, the popliteal node (pLN). The ability to differentiate and quantify the very high signal in the footpad from the low signal in the lymph node is modeled using samples of SPIO and injections of SPIO-labeled DC in mice.

## Methods

### Cell Labeling

#### Animals

C57BL/6 mice were sourced from the breeding operation at the West Valley Barrier Facility at Western University. All applicable animal protocols were approved by the University of Western Ontario Animal Care and Use Subcommittee.

#### Mouse bone marrow-derived dendritic cell (DC) culture

Mouse bone marrow-derived DC were generated from the femurs and tibias of C57BL/6 mice and cultured as previously described by our group and others.^18–20^ Briefly, bone marrow progenitor cells were cultured in complete RPMI media supplemented with interleukin-4 (IL-4, 10 ng/mL, PeproTech, Canada) and granulocyte-macrophage colony-stimulating factor (GM-CSF, 4 ng/mL, PeproTech, Canada) for four days. On day 4, immature DC were enriched from culture using Histodenz™ (Millipore Sigma, Canada) gradient centrifugation.

#### SPIO labeling of DC

Following gradient enrichment of immature DC, DC were labeled with Synomag-D (Micromod GmbH) (200 μg Fe/mL) added to culture via simple co-incubation in complete RPMI (+IL-4 and GM-CSF). To increase Synomag-D uptake by DC, transfection agents (TAs) were employed. Protamine sulfate (0.24 mg/mL) and heparin (8 USP units/mL) were individually diluted in OPTI-MEM in Eppendorf tubes. Following Synomag-D addition to OPTI-MEM containing heparin, the contents of both Eppendorf tubes were combined, vortexed and added to immature DC. Synomag-D was added such that DC were cultured in a final concentration of 200 μg Fe/mL for 4 hours, after which time complete RPMI (+IL-4 and GM-CSF) was added to ensure DC were cultured overnight at the same concentration as DC labeled via simple co-incubation. On day 5 of culture, regardless of which SPIO labeling method was used on day 4, a previously defined maturation cocktail^21^ was added to DC which were then cultured for an additional 24 hours.

#### Magnetic Selection of SPIO+ DC

To ensure that only Synomag-D+ DC were adoptively transferred, magnetic column enrichment was performed as previously described.^22^ Briefly, on day 6 mature DC were collected, extensively washed in PBS and then resuspended in 2 mL PBS per DC culture condition in a 12 × 75 mm sterile tube. DC suspensions were incubated for five minutes in an EasySep^™^ magnet (Stemcell Technologies, Vancouver, CAN) at room temperature. After five minutes, both tube and magnet were inverted to collect flow through containing unlabeled DC while Synomag-D+ DC remained in the tube and were collected, counted and prepared for subsequent adoptive cell transfer. An aliquot of Synomag-D+ DC was collected to histologically confirm Synomag-D uptake through Perls Prussian Blue (PPB) staining. Unlabeled DC were used for viability and immunophenotyping analysis.

### Flow Cytometry

DC were phenotyped and assessed for viability upon completion of a 6 day culture using a previously outlined protocol.^23^ Briefly, Zombie NIR™ fixable viability dye (Biolegend, San Diego, USA) was employed to assess DC viability and preceded surface immunofluorescence staining. In the presence of TruStain FcX™ anti-mouse CD16/32 block (clone: 93), DC were stained with PE CD45 (30-F11) and APC CD11c (N418) for 25 minutes at 4°C (all Biolegend). Excess antibodies were removed by washing with HBSS + 0.1% BSA. Upon resuspension in HBSS + 0.1% BSA, DC were fixed with 4% paraformaldehyde and kept at 4°C until acquisition using a LSRII analytical flow cytometer (BD Biosciences, San Jose, USA). FlowJo software (v10, Tree Star, Inc., Ashland, USA) was used for all analyses.

### *In vitro* MPI

#### MPI Relaxometry

To evaluate the sensitivity and resolution of Synomag-D it was compared to another SPIO called Vivotrax (Magnetic Insight Inc.) Vivotrax is a ferucarbotran previously used as a liver MRI contrast agent under the brand name Resovist^®^. It was adopted early for MPI and is one of the most common SPIO used now for MPI. Using the Relax^™^ module equipped on the Momentum^™^ scanner, measurements were collected in triplicate for: (i) Vivotrax (ii) Synomag-D (iii) Synomag-D with TAs (iv) Synomag-D+ DC with TAs.

For (i) and (ii), 1 μL of the agent was used. For (iii) 1.25% of the volumes used for the labeling procedure were prepared (1.125μL Synomag-D, 62.5μL serum free media, 0.0375μL protamine sulfate, 0.0125μL heparin). For (iv), 500K labeled DC suspended in 40 μL PBS were collected in a PCR tube. Point spread functions (PSF) from MPI relaxometry were analyzed for sensitivity (peak signal) and resolution (full width half maximum, FWHM), with signal normalized to the iron content of each sample. To compare resolutions, FWHM values reported in milli-Tesla (mT) were converted to millimetres (mm) by dividing by the 3.0 Tesla per meter (T/m) gradient strength. Ordinary one-way ANOVA tests were performed to determine differences in sensitivity and resolution for relaxometry measurements.

#### DC detection with MPI

After Synomag-D+ DC magnetic column enrichment, triplicate sets of cell samples containing 250K, 100K, 50K, 25K, 12K, 6K, (K = thousand) DC labeled with TAs (n = 18) and without TAs (n = 18) in 40 μL PBS were prepared and imaged with MPI. Cellular sensitivities were estimated for both labeling methods by graphing iron content versus number of DC and comparing linearly fitted slopes. All images were acquired on a Momentum^™^ MPI scanner (Magnetic Insight Inc.) using 2D high sensitivity isotropic (3.0 T/m gradient) imaging and a 6 × 6 cm field of view (FOV). A simple linear regression was performed to determine the relationship between iron content and cell number.

#### DC migration model

To model DC migration, a 250K cell sample (high signal) was placed 2 cm from a 25K cell sample (low signal) to resemble the distance and signal between the injection site (footpad) and the pLN. The 25K cell sample was then moved incrementally 1 cm away, up to 5 cm, from the 250K cell sample to determine the minimum distance at which two distinct signals could be resolved. Images were acquired in 2D and 3D using high sensitivity isotropic (3.0 T/m gradient) parameters and a 12 × 6 cm FOV.

### *In vivo* MPI

Four *in vivo* experiments were performed: (i) DC migration model (following *in vitro* modelling experiments), (ii) a mock DC migration experiment, (iii) a pilot DC migration experiment, and (iv) a true migration experiment. Mice were anesthetized initially with 2% isoflurane and maintained 1% isoflurane during imaging. All images were acquired in 2D and 3D using high sensitivity isotropic (3.0 T/m gradient) parameters and a 12 × 6 cm FOV.

For the DC migration model (i), 25K and 250K Synomag-D+ DC in 40 μL PBS were subcutaneously injected into the upper back of anesthetized C57BL/6 mice (n = 4) at distances of 2 cm (n = 2) or 5 cm (n = 2) apart. MPI was immediately performed post-injection.

In the mock DC migration (ii) 6K Synomag-D+ DC were adoptively transferred into the left and right popliteal lymph node region of a C57BL/6 mouse (n = 1). Assuming a migration of ~5%, 125K Synomag-D+ DC were subcutaneously injected into the ipsilateral hind left and right footpads of the same mouse. All injections were formulated in 40 μL PBS. MPI was immediately conducted post-injection. In this experiment, 6K DC was chosen to repeat the number of DC detected *in vitro*.

For the pilot migration (iii) 300K cells were injected into each hind footpad and imaging was conducted the same day (day 0) and repeated 48 hours later (day 2) to detect signal in the pLNs from migratory DC. The left and right hind limbs (footpad and pLN) were removed after fixation by cardiac perfusion and imaged *ex vivo*. Further validation of pLN signal was done by imaging excised lymph nodes. Iron mass was quantified from measured MPI signal, as described below.

For the true *in vivo* migration experiment (iv) 300K (n = 3) or 500K (n = 3) Synomag-D+ DC were intradermally injected into each hind footpad immediately following Tag-It Violet^™^ (Violet+) cell tracking dye incorporation (Biolegend, San Diego, USA). MPI was done immediately following injection (day 0) and repeated 48 hours later (day 2). For this experiment, mice were individually fasted overnight for 12 hours prior to imaging with only water, a laxative, and corn bedding. After imaging, mice were returned to fully supplied cages with food. This was done to reduce gastrointestinal signal in MPI images. On day 3 whole mouse body proton MRI was acquired on two of the mice that showed the highest MPI pLN signal from each injection group using a clinical 3T MRI (Discovery MR750, General Electric) system and a 4.3 × 4.3 cm surface coil (Clinical MR Solutions, Wisconsin) as described.^24^ Left and right pLNs from all mice were excised and imaged *ex vivo* with MPI on day 3. pLNs were prepared and sectioned as previously described.^19^ Qualitative imaging of pLNs was performed to identify Violet+ Synomag-D+ DC using an Olympus IX50 phase contrast inverted microscope (Richmond Hill, CA) and Infinity3-3URF camera (Lumenera, Ottawa, CA). Iron mass was quantified from measured MPI signal, as described below. Additionally, two cell pellet samples each containing 1000K DC were imaged using 2D and 3D high sensitivity isotropic (3.0 T/m gradient) parameters to measure the amount of iron per cell and subsequently estimate cell number.

### Quantification

Calibration lines (2D and 3D) for Synomag-D were made to determine the relationship between iron content and MPI signal using previously established methods.^24^ All MPI images were imported and viewed with a custom MPI colour look-up table (CLUT) using the open-source Horos™ image analysis software (version 3.3.6, Annapolis, MD USA). MPI signal was measured within a specific region of interest (ROI) using a semi-automatic segmentation tool for both 2D and 3D images. For each ROI the window/level (W/L) was set to the minimum and maximum signal, displaying the full dynamic range for that region. ROIs were extended to half the maximum signal value of that region to ensure real signal above background levels was measured. Total MPI signal for an ROI was calculated by multiplying the ROI area (2D) or volume (3D) by the mean signal. Iron content was calculated by dividing the total MPI signal by the slope of the calibration line. All MPI images (calibrations, *in vitro* DC pellets, and *in vivo* experiments) were delineated and analyzed in the same way to ensure consistency. Iron content per cell was calculated by dividing the measured iron content from the signal ROI by the number of cells in the cell pellet sample (1000K). Cell numbers for the true migration experiment were then estimated by dividing the iron quantified within a given ROI by the respective iron/cell measurements. Unpaired t-tests of the means were used to compare *in vivo* and *ex vivo* quantification.

## Results

### *In vitro* MPI

#### MPI Relaxometry

The relaxometer mode on the MPI system was used to compare the performance of the SPIOs Synomag-D and Vivotrax and to determine if there are differences after Synomag-D is internalized by cells. Figures 1A and B show the point spread functions (PSF) which provide information on the relative sensitivities and resolutions, for the free SPIOs, Synomag-D mixed with TAs and Synomag-D in cells. The resolution in mm was determined by dividing the FWHM of the PSF by the 3.0 T/m gradient strength. Synomag-D had approximately 3.46 times higher sensitivity and improved resolution (2.49 mm versus 3.29 mm) compared to Vivotrax. This agrees with Vogel et al.^25^ Figures 1D and E summarize the results comparing free Synomag-D, Synomag-D mixed with TAs and intracellular Synomag-D. Sensitivity was reduced when Synomag-D was mixed with heparin and protamine sulfate and was further reduced when internalized into cells. There was no statistically significant difference in resolution between Synomag-D and Synomag-D with TAs, however, the resolution was significantly lower for intracellular Synomag-D (Figure 1E). The negative effect on the MPI signal which occurs with mixing with TAs and internalization has been shown previously and is the result of aggregation and the slowed MPI relaxation.^26^

**Figure 1:**
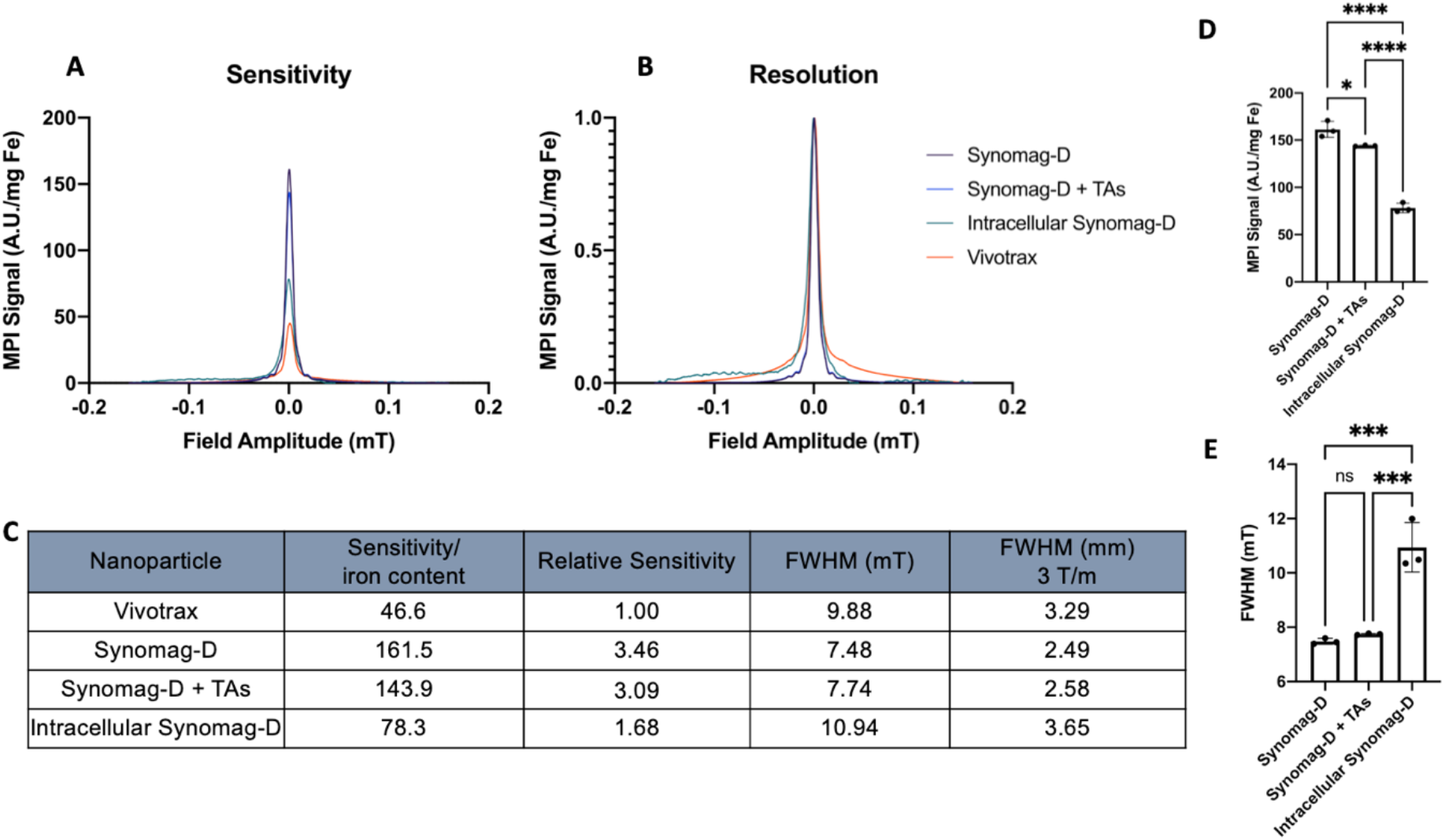
Comparing MPI sensitivity and resolution of Vivotrax, Synomag-D, Synomag-D with transfection agents (TAs), and intracellular Synomag-D. Relative sensitivities are compared in (A) with MPI signal normalized to iron content of the sample. Relative resolutions are compared in (B) with MPI signal normalized to iron and maximum sensitivity values. Numerical values for the graphs are shown in (C). Highly significant differences were seen in both sensitivity (D) and resolution (E) with intracellular synomag-D. There was a significant difference in sensitivity (D) and a non-significant difference in resolution (E) between Synomag-D and Synomag-D with TAs.

#### DC detection with MPI

PPB staining showed the presence of iron in DC labeled with TAs (Figure 2A) and without TAs (Figure 2B). The addition of TAs improved DC labeling with both a higher amount of iron/cell (5.5 pg Fe/cell vs 1.4 pg Fe/cells) and more iron positive cells (76.6% vs 51.6%). Cell viability was not affected by cell labeling compared to unlabeled cells (Figure 2C). Both labeling strategies show strong linear correlations between measured iron content and cell number, as expected (Figure 2D). DC labeled with TAs had a higher cellular sensitivity than no TAs, indicated by a steeper slope (5.73 × 10^−6^ vs 1.24 × 10^−6^). MPI of Synomag-D labeled DC phantoms without TAs is limited to a detection of 25K cells due to lower labeling efficiency (Figure 2E). With more iron/cell from enhanced labeling with TAs, 6K cells are visible with MPI, the lowest number tested in this experiment (Figure 2F). The addition of TAs improved cell labelling compared to no TAs, confirmed by PPB stains, MPI, and iron quantification. This was the labeling strategy used for all further experiments.

**Figure 2:**
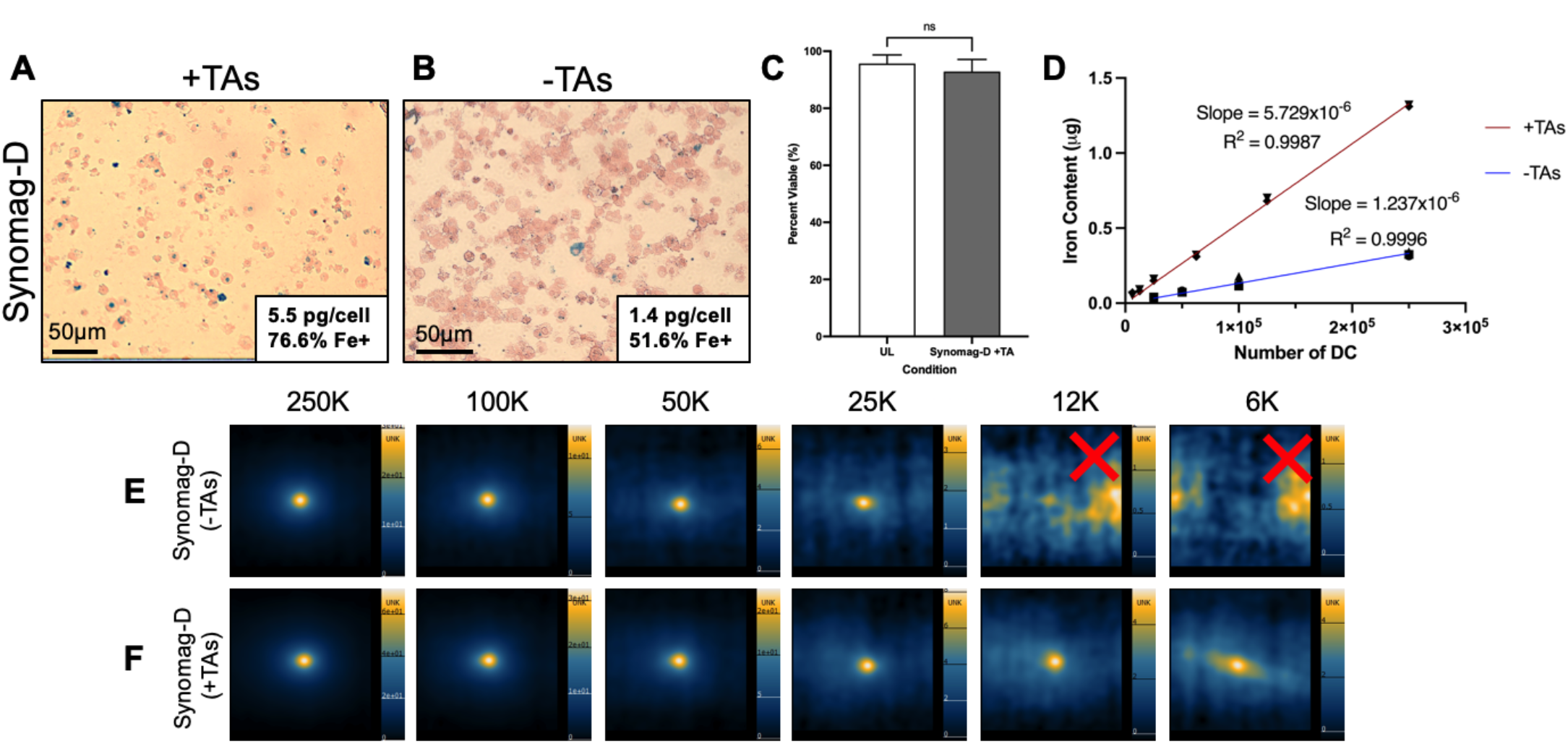
DC labeling and MPI cellular detection with Synomag-D. PPB staining shows the presence of iron (Synomag-D) in DC labeled with TAs (A) and without TAs (B). The addition of TAs showed a higher labeling efficiency (5.5 pg/cell vs 1.4 pg/cell) and more Synomag-D+ cells (51.8 % vs 76.6 %). CD11c+ DC viability is unaffected by Synomag-D labeling or the use of TAs (C). Flow cytometry was used to identify singlet, CD45+CD11c+ DC (gating not shown) and subsequently assessed for viability using Zombie NIR^™^ fixability viability dye. CD11c+ DC viability was unchanged (grey bar) following Synomag-D labeling (200 μg/mL) in the presence of TAs (protamine sulfate and heparin) when compared to unlabeled CD11c+ DC from the same culture (white bar). Data shown as mean ± SD (n = 4, ns - no statistical significance, p > 0.05, paired t-test). MPI cellular sensitivities are shown in (D). DC labeled with TAs have a higher cellular sensitivity than without TAs, indicated by a steeper slope (5.729 × 10^−6^ vs 1.237 × 10^−6^). Iron content was quantified from MPI images of cells (E, F). Detection was limited to 25K cells without using TAs, below which signal was not detected, indicated by the red X (E). With TAs, 6K cells are visible, the lowest number tested in this experiment (F).

#### In vitro modelling of DC migration

Using 2D imaging, samples with equal numbers of Synomag-D+ DC can be clearly discerned when placed 2 cm apart. Two samples of 25K cells (Figure 3A) and 250K cells (Figure 3B) placed 2 cm apart are equally visible with two distinct peaks shown in the adjacent corresponding signal intensity profile. When the 25K cell phantom is imaged with the 250K cell phantom, creating unequal signals, the signal from fewer cells (less iron) was obscured by the signal from more labeled cells. The corresponding signal intensity profile shows the 25K cell phantom as a small shoulder off the tail of the higher peak from the 250K cell phantom (Figure 3C). To fully separate the unequal signals into two distinct peaks, a minimum distance of 5 cm was required for 2D imaging (Figure 3D). Windowing the image to the minimum and maximum signal from the 25K cell phantom oversaturated MPI signal from the 250K cell phantom, expanding into the region of the lower signal, and prevented detection of the 25K cell phantom. Distinct signals were visible using 3D imaging at 2 cm, however window leveling to the 25K cell phantom signal still oversaturated signal from the 250K cell phantom (Figure 3E).

**Figure 3:**
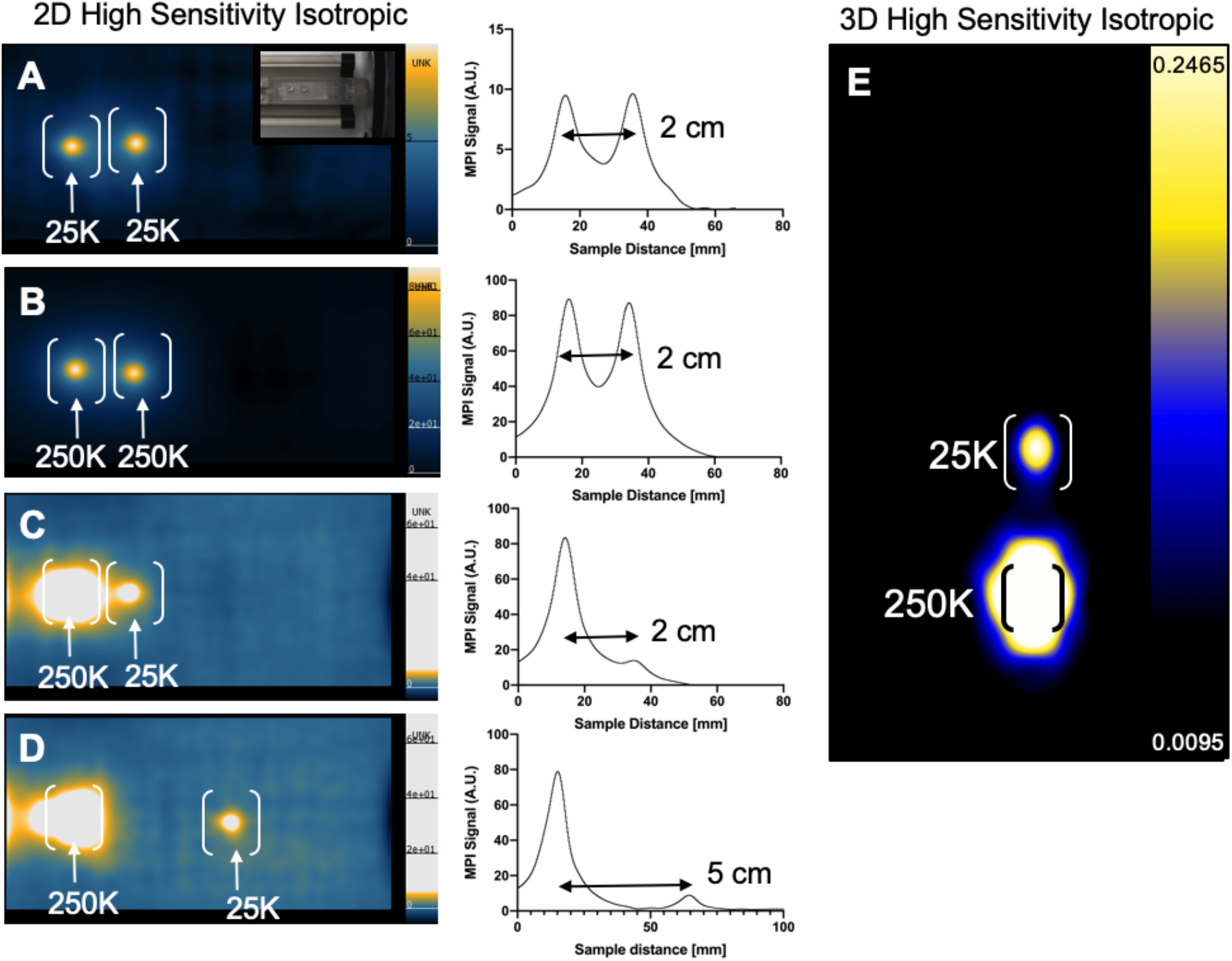
Imaging two sources of unequal MPI signal within close proximity to each other. Two equal samples of either 25K (A) or 250K (B) Synomag-D+ DC are clearly visible and can be resolved at 2 cm separation (images shown in full dynamic range). Signal from 25K cells becomes partially hidden when placed 2 cm from 250K cells (C). At a distance of 5 cm, both signals could be fully resolved with 2D imaging (D). Window leveling for images (C) and (D) is set to the minimum and maximum signal of the 25K cell sample. Corresponding signal intensity profiles are shown adjacent to images. With 3D imaging, (as in the samples and position in C) both signals can be resolved at 2 cm (E). Data is displayed in landscape mode for 2D images and portrait mode for 3D.

### *In vivo* MPI

In experiment (i) the *in vitro* DC migration model was repeated to compare signal detection and resolution between two unequal signal sources in an *in vivo* mouse model. As with the *in vitro* model, subcutaneous injections in mice of 25K and 250K cells 2 cm apart (Figure 4A) could not be resolved with 2D imaging (Figure 4B) but could be with 3D imaging (Figure 4C). To clearly see the 25K cell injection site the contrast and intensity of the images had to be adjusted (4D). When imaging mice with cell injections 5 cm apart (Figure 4E) gastrointestinal (GI) signal was detected in all mice, coming from iron contained within mouse feed and ingested cage bedding (Supplementary Figure 1). Three distinct signal regions were visible with both 2D and 3D imaging (i) 250K cells, (ii) GI, and (iii) 25K cells. Signal from the lower source (25K) is partially hidden within signal from the GI using 2D imaging (Figure 4F). With 3D imaging, signal from 25k is not visible with the image shown in full dynamic range (Figure 4G). However, after adjusting the window/level to the signal from the 25K cell injection, a distinct ROI is clearly visible (Figure 4H).

**Figure 4:**
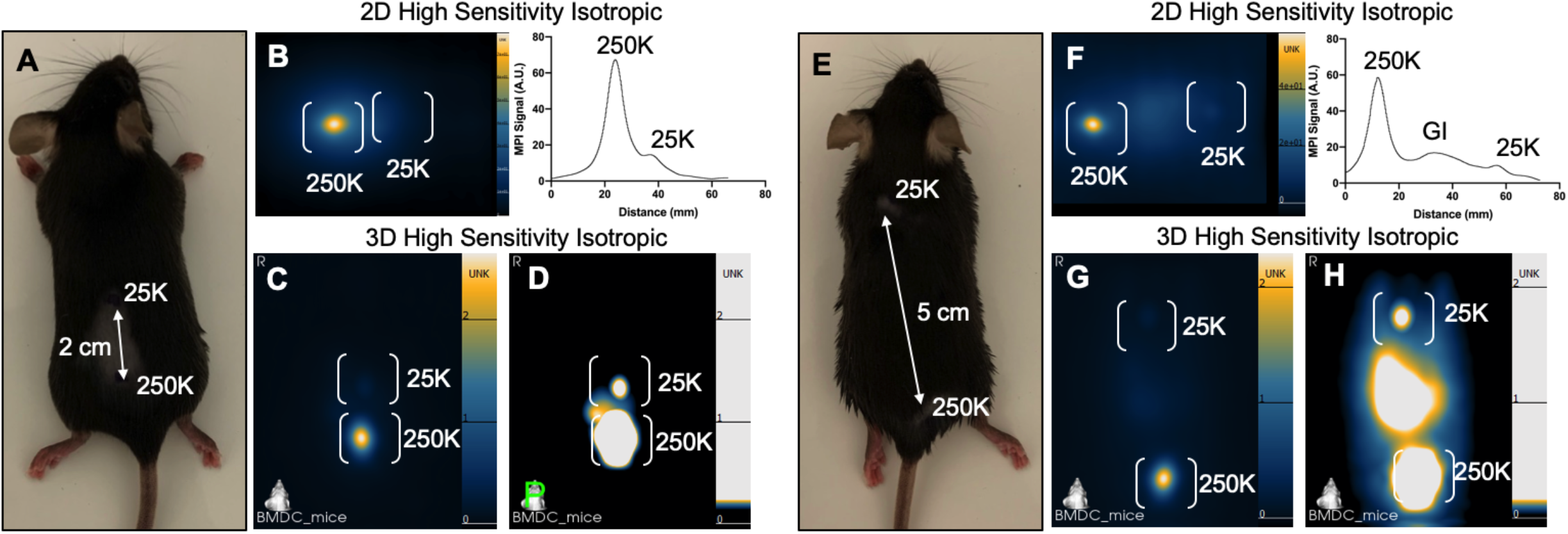
*In vivo* cellular sensitivity and resolution after 25K and 250K Synomag-D+ DC were injected subcutaneously into the upper back of anesthetized C57BL/6 mice (n = 4) separated by either 2 cm (n = 2) or 5 cm (n = 2). At a distance of 2 cm (A), in 2D images, the signal from 25K cells is obscured by the signal from 250K cells, with unresolved peaks in the corresponding signal intensity profile (B). With 3D imaging (C, D) the two regions of signal can be resolved after window leveling to the lower signal (D). When separated by 5 cm (E) the signal from the 25K cells is partially obscured by stronger signal from the GI tract (F), shown in the corresponding signal intensity profile. With 3D imaging (G, H) signal from 25K cells is resolved after window leveling to the 25K cell sample (H). Data is displayed in landscape mode for 2D images and portrait mode for 3D.

In experiment (ii) a mock migration was performed on a C57BL/6 mouse (n = 1) to mimic the expected signals coming from the footpads and the pLNs with 6K Synomag-D+ DC injected into the left and right pLNs and 125K Synomag-D+ DC injected into the left and right footpads (Figure 5A). Our choice of 6K for this experiment was based on our *in vitro* findings. With 3D imaging, signal was clearly visible at the injection sites of the left and right pLNs, as well as signal from the footpads and strong GI signal (Figure 5B). *In vivo* signal from 6K cells had an estimated iron content of 0.010 μg Fe (left) and 0.017 μg Fe (right) and signal from 125K cells in the footpads had an estimated iron content of 0.104 μg Fe (left) and 0.132μg Fe (right).

**Figure 5:**
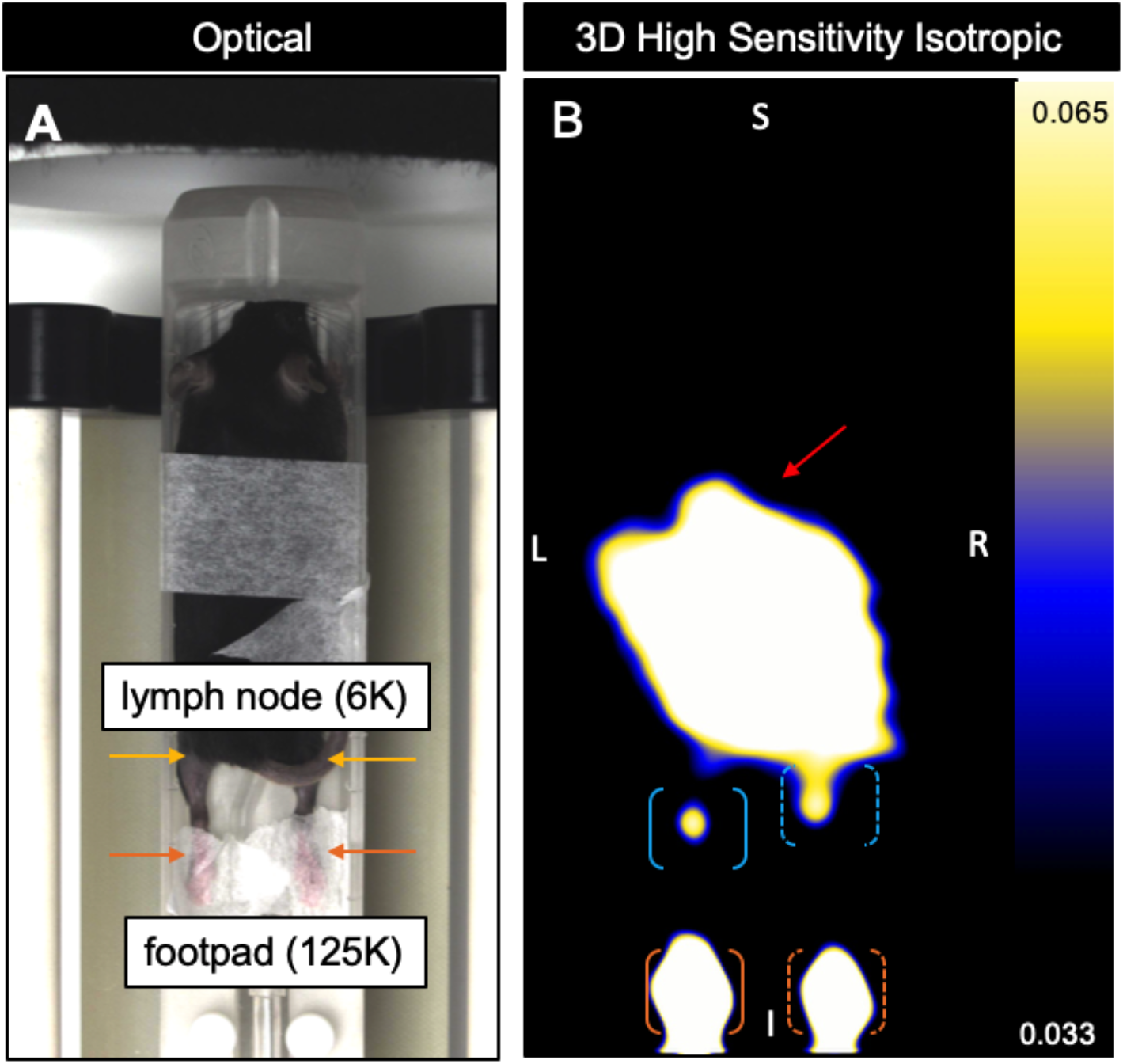
Mock migration experiment mimicking the expected distance and signal using Synomag-D+ DC injected subcutaneously into the left and right pLN (6K, yellow arrows) and footpad (125K, orange arrows) (A). MPI was acquired using high sensitivity isotropic (3.0 T/m gradient) parameters. In a slice selected from 3D imaging (B) signal from the left (blue bracket) and right (blue dashed bracket) pLNs are visible and quantifiable. Window leveling to the pLN signal oversaturates signal from the GI tract (red arrow) and footpads (left - orange bracket, right - orange dashed bracket). Iron quantification is as follows: left pLN = 0.010 μg Fe, right pLN = 0.017 μg Fe, left footpad = 0.104 μg Fe, right footpad = 0.132 μg Fe.

In experiment (iii) a pilot DC migration experiment was performed on a C57BL/6 mouse (n = 1) to test the ability of MPI to detect migratory DC to the pLNs following the injection of 300K, a biologically relevant number.^19^ For this experiment, only iron was quantified because the iron content/cell was not measured from a cell pellet sample to estimate cell numbers. Day 0 MPI only showed signal in the left and right footpads, as expected, with 0.282 μg Fe in the left and 0.352 μg Fe in the right footpad (Figure 6A). On day 2 MPI signal was detected in the pLNs, indicating DC migration (Figure 6B). Windowing and leveling to the minimum and maximum signal of the pLNs showed a defined, clear signal region with 0.050 μg Fe in the left pLN (Figure 6C) and 0.013 μg Fe in the right pLN (Figure 6D). The MPI signal in the footpads was reduced on day 2, as expected, also suggesting DC migration with an estimated 0.208 μg Fe in the left footpad and 0.128 μg Fe in the right footpad. The hind limbs were removed to allow for validation of the *in vivo* findings. Clear MPI signal was detected in both footpads and pLNs in the *ex vivo* hind limb images, confirming the presence of iron in the lymph nodes. Iron content was measured with 0.058 μg Fe in the left pLN and 0.254 μg Fe in the left footpad (Figure 6E) and 0.021 μg Fe in the right pLN and 0.176 μg Fe in the right footpad (Figure 6F). Excised pLNs showed clear MPI signal with 0.077 μg Fe in the left pLN (Figure 6G) and 0.013 μg Fe in the right pLN (Figure 6H).

**Figure 6:**
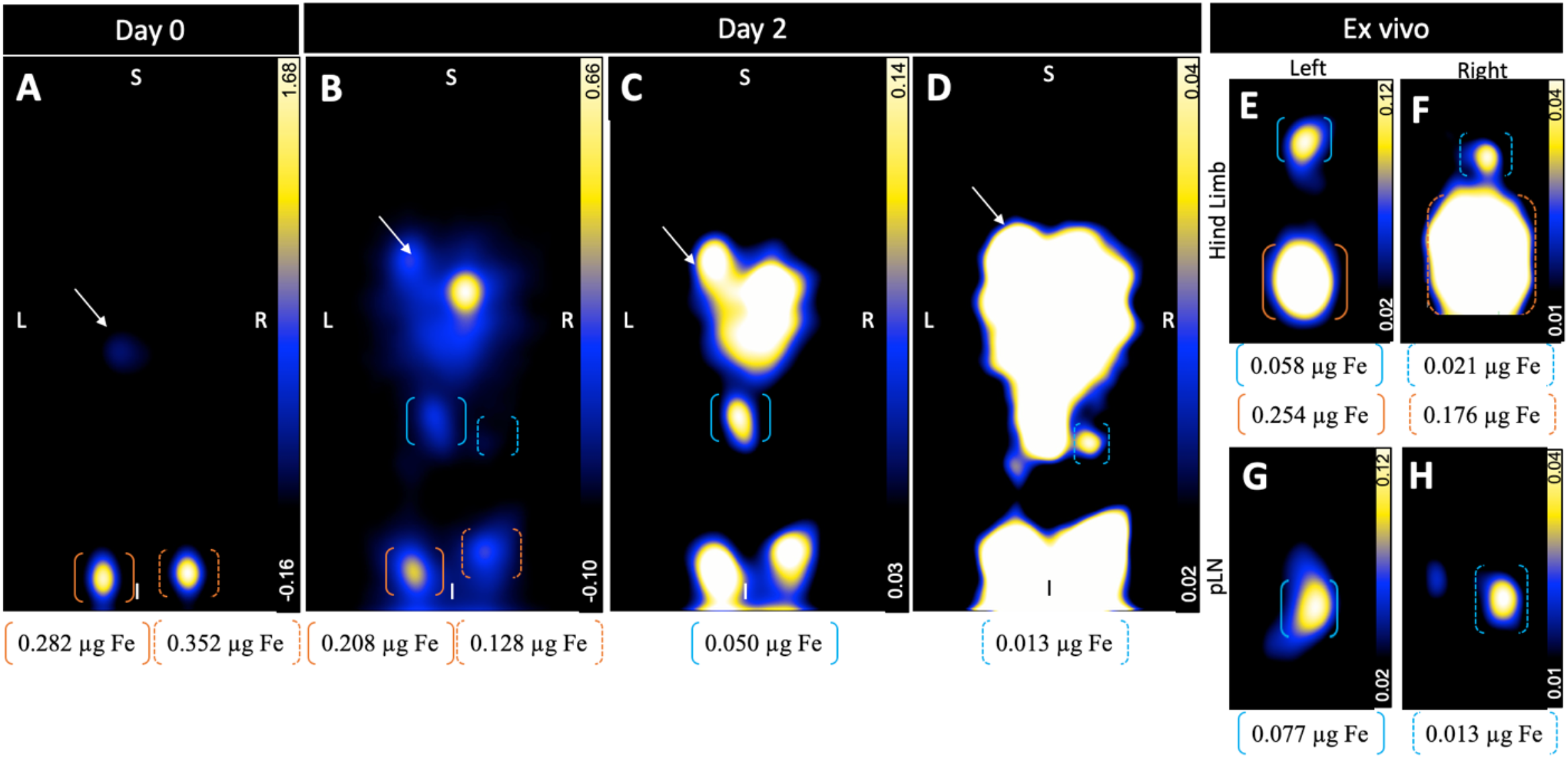
Pilot migration experiment showing *in vivo* and *ex vivo* MPI of C57BL/6 mouse DC migration following injection of 300K Synomag-D+ DC into the hind left and right footpads. All images are slices selected from 3D high sensitivity isotropic (3.0 T/m gradient) images with minimum and maximum MPI signal values indicated by the scale bars. Day 0 imaging (A) displays two signal regions in the hind footpads (left - orange brackets, right - orange dashed brackets). Day 2 imaging in (B) suggests DC migration to the pLNs (image shown in full dynamic range) with clearer images window leveled to the minimum and maximum signal of the left pLN (blue bracket, C) and right pLN (blue dashed bracket, D). Oversaturation of GI (A-D, white arrow) and footpad signal is increasingly apparent with extreme window leveling to the lower signal intensities from the pLNs. Hind limbs were imaged *ex vivo* and show clear signal in the left (E) and right (F) footpad and pLN. Excised pLNs were imaged confirming DC migration in both the left (G) and right (H) pLNs. Iron quantification is shown underneath for the corresponding brackets in respective images.

In experiment (iv) a true migration experiment was performed in mice (n = 6) injected with either 300K (n = 3) or 500K (n = 3) Violet+ Synomag-D+ DC. In this experiment mice were fasted which effectively removed GI signal so that it was not visible within the dynamic range of the pLN signals. Table 1 summarizes the data from all 6 mice, showing cell number and iron estimates for the footpads and pLNs on day 0, day 2, and *ex vivo*. Footpad signal (left and right) was clearly visible and quantifiable on both day 0 and day 2 imaging. On day 2, four of the six mice showed MPI signal in the left pLN and two of the six mice showed MPI signal in the right pLN. The number of DC injected did not affect the ability to detect MPI signal in the pLNs; for each cell number, two of the three mice had signal in the left pLN, and one of the three mice had signal in the right pLN. *Ex vivo* imaging of the pLNs improved detection of MPI signal; signal was detected in 11/12 nodes. There was no significant difference, however, in the mean values for iron content or cell number for *in vivo* versus *ex vivo* measurements (Figure 7).

**Table 1:**
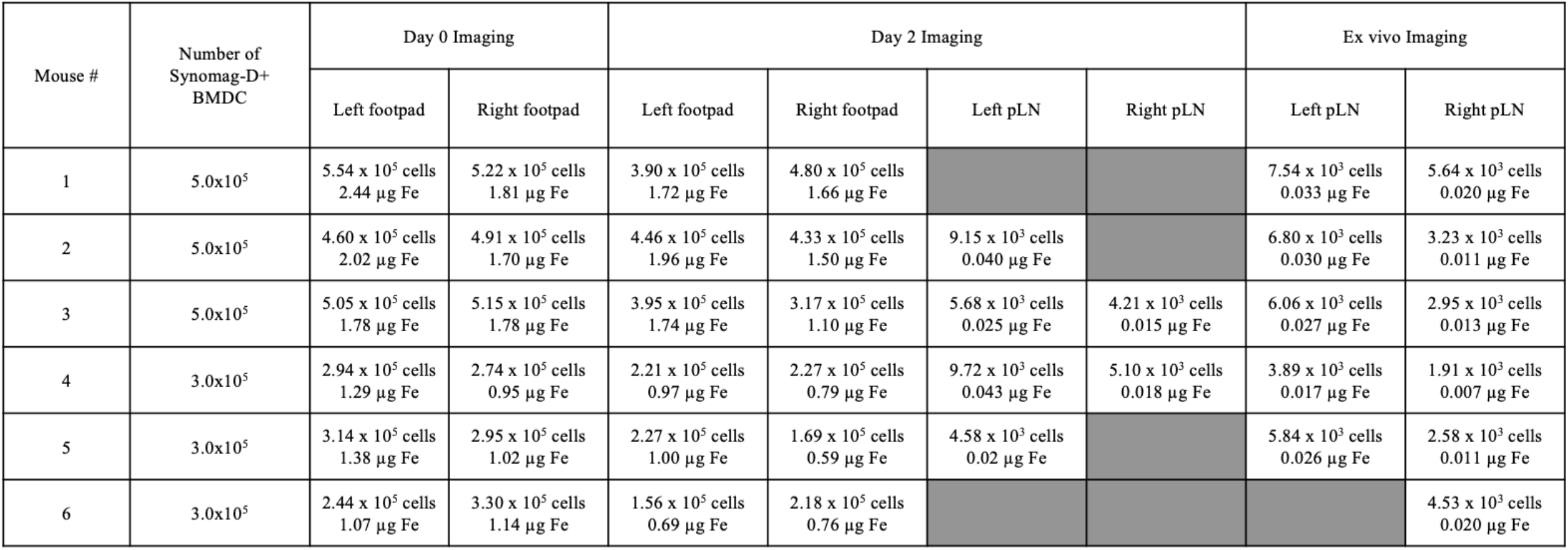
Quantification for the true migration experiment (iv) in C57BL/6 mice (n = 6) showing cell number estimates with respective iron content (μg) for Day 0, Day 2, and *ex vivo* imaging. Grey squares indicate undetectable signal.

**Figure 7:**
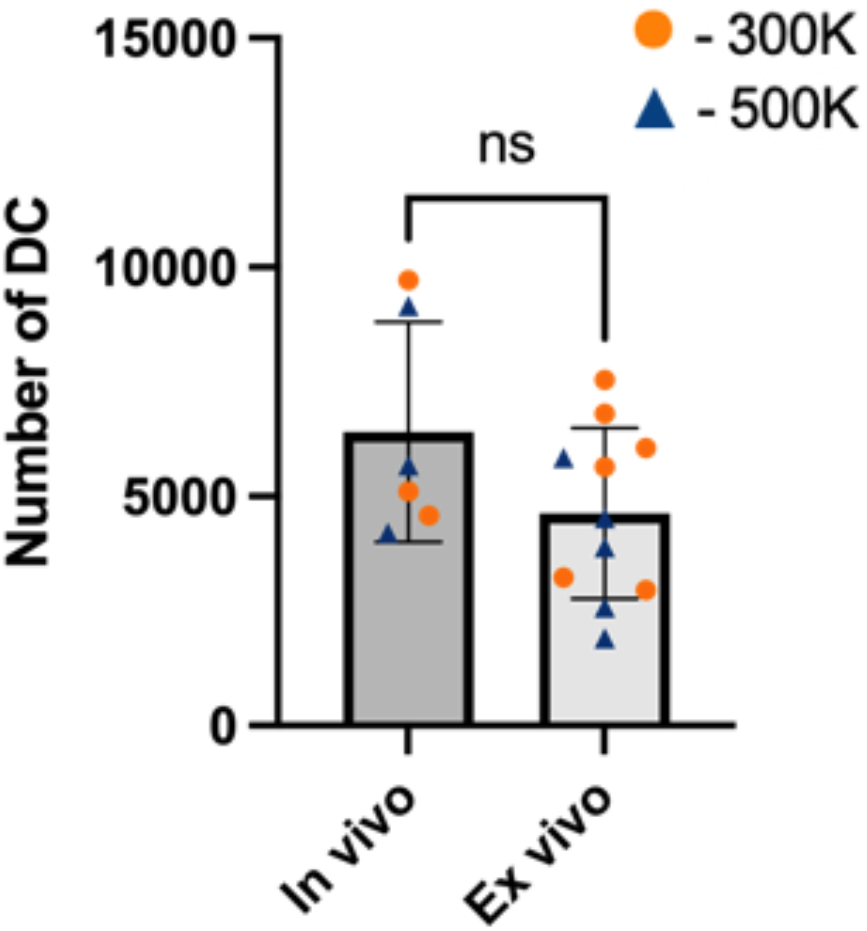
Quantification statistics for the true migration experiment including the 300K and 500k injection groups (iv). There was a non-significant difference between *in vivo* and *ex vivo* quantification. Data shown as mean ± SD (ns - no statistical significance (p > 0.05), unpaired t-test).

Figure 8 shows images from one representative mouse (mouse 4) imaged with both MPI and MRI, with 300K cells injected into the footpads. In this mouse, MPI signal was detected in both the left and right hind footpads on day 0, as expected, with an estimated 294K cells (1.29 μg Fe) in the left footpad and 274K cells (0.95 μg Fe) in the right footpad (Figure 8A). Day 2 MPI showed signal in the left and right pLNs, with approximately 9720 cells (0.043 μg Fe) in the left pLN and 5100 cells (0.018 μg Fe) in the right pLN, suggesting a 3% and 2% migration rate, respectively (Figure 8B). Window leveling was required to visualize pLN signal compared to the stronger footpad signals. Day 3 MRI showed regions of signal loss in both the left and right pLNs (Figure 8C). This was also true for the second mouse imaged with both MPI and MRI (mouse 3, not shown). The signal void volumes were 0.31 mm^3^ (left) and 0.35 mm^3^ (right) for mouse 3 and 0.43 mm^3^ (left) and 0.32 mm^3^ (right) for mouse 4. Representative *ex vivo* MPI of the left and right pLNs is shown in Figure 8D for mouse 4. MPI of pLNs excised on day 3 (after MRI) showed clear signal in both pLNs, with an estimated 3890 cells (0.017 μg Fe) in the left pLN and 1910 cells (0.007 μg Fe) in the right pLN. After imaging with MPI and MRI, left and right pLNs were removed from all mice and sectioned for fluorescence imaging. Figure 9 shows representative images of sectioned pLNs. Violet+ DC appear centrally in the nodes, as expected for pLN migrated DC, in both left (Figure 9A) and right (Figure 9B) pLNs.

**Figure 8:**
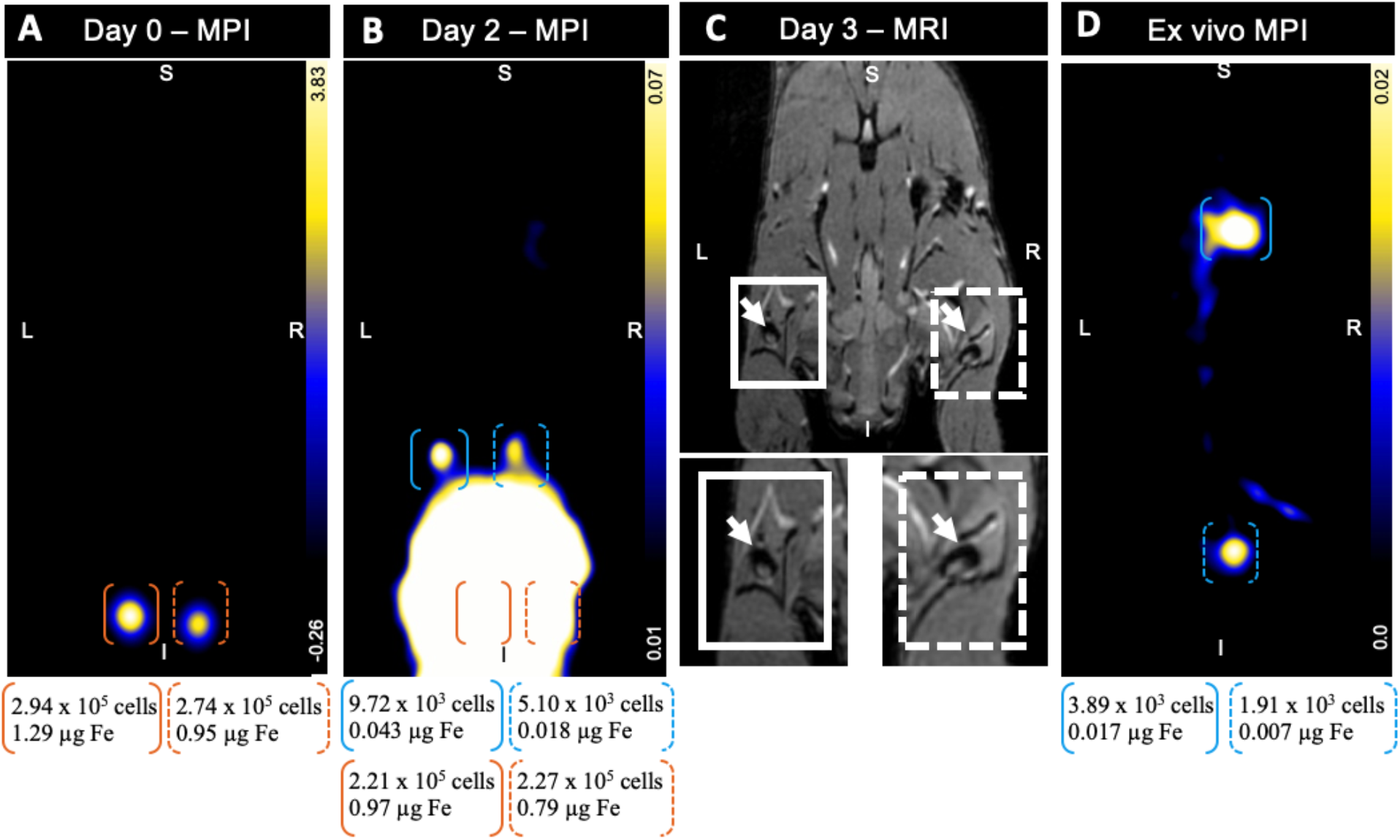
True DC migration experiment (iv) showing *in vivo* and *ex vivo* MPI of C57BL/6 mouse DC migration following injection of 300K Synomag-D+ DC into the hind footpads. All images are slices selected from 3D high sensitivity isotropic (3.0 T/m gradient) images. MPI signal values are indicated by the scale bars. Day 0 imaging (A) displays two regions of MPI signal from the hind footpads (left - orange bracket, right - orange dashed bracket). Day 2 imaging (B) shows signal in both pLNs (left - blue bracket, right - blue dashed bracket). Oversaturation of footpad signal is apparent with extreme window leveling to the lower signal intensities of the pLNs. On day 3 coronal MRI (C) shows clear signal loss in the pLNs (left - white box, right - white dashed box) indicated by the white arrows. *Ex vivo* MPI of excised pLNs shows clear signal in the left and right pLNs, also confirming DC migration (D). Iron quantification and cell number estimates are shown below images.

**Figure 9:**
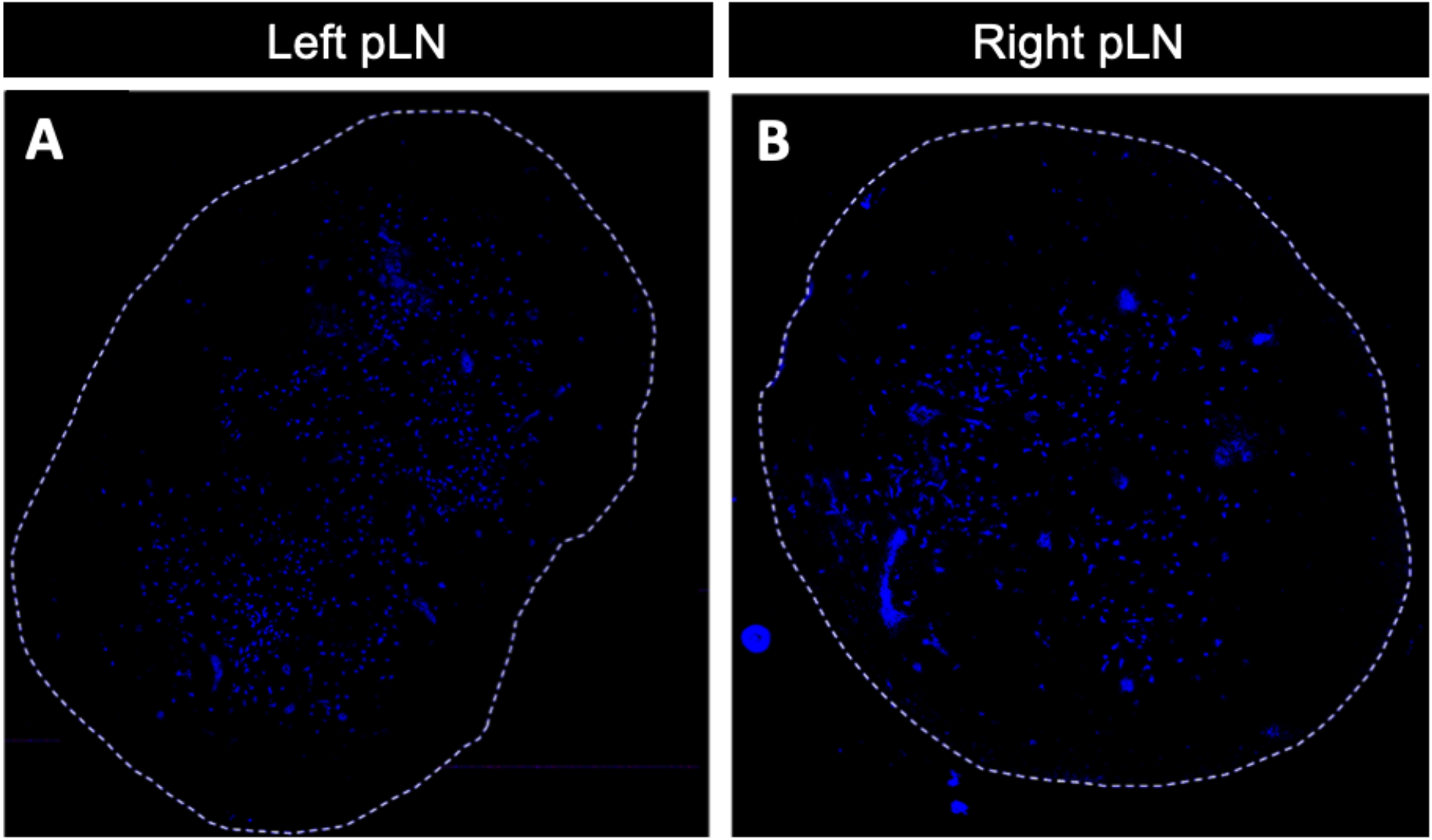
Qualitative fluorescent images of pLN cryosections from mouse 3 revealed migrated Violet+ Synomag-D+ DC in both the left (A) and right (B) pLNs.

## Discussion

This study demonstrates the excellent potential for MPI as a non-invasive technique to detect and quantify the migration of DC to lymph nodes. In the regime of cancer vaccination DC can generate effective and long-lasting immune responses. The recruitment of DCs to lymph nodes is crucial for the induction of immune responses and the success of DC immunotherapy is strongly related to the number of administered DC that accumulate in the nodes.^27,28^ However, the best route and timing for DC administration and the ideal strategies for maturation and activation of DC to maximize migration and accumulation are not clear. *In vivo* imaging can be used to verify the accuracy of the DC injection, to image the spatial distribution and migration of DC after administration and is being investigated as a biomarker for evaluating migration efficacy.

Iron-based MRI cell tracking has been the most widely used imaging modality for preclinical studies of DC migration.^18,21,29,30^ Dekaban et al. and Bulte et al. have reviewed this.^31,32^ Experimental studies in mice have demonstrated that SPIO-labeled DC can be detected as regions of signal loss in T2- or T2*-weighted images after their intradermal or intranodal administration. These studies used measurements such as signal void volume, number of black pixels, signal intensity or fractional signal loss to relate the signal loss to the number of cells injected, histological analysis of DC in nodes, the iron content in lymph nodes measured *ex vivo* or tumor size.^18,21,22,30,33–37^ De Chickera et al. showed that MRI could detect differences in the migratory abilities of two different DC preparations, *ex vivo* untreated, resting DC versus *ex vivo* matured DC; both the signal void volume and the fractional signal loss were significantly increased following addition of a cytokine maturation cocktail and the toll-like receptor 9 ligand CpG.^21^ Grippin et al. showed that T2*-weighted MRI intensity in lymph nodes is a strong correlate of DC trafficking and an early predictor of antitumor immune response.^38^ Relative signal intensity was calculated as the MRI intensity of the treated lymph node divided by the MRI intensity of the untreated lymph node. Two days post vaccination tumor-bearing mice with high relative signal intensities in the draining lymph nodes had significantly smaller tumors than mice with low relative signal intensities. Despite the evidence that preclinical MRI of SPIO-labeled DC can be used to assess the efficacy of DC immunotherapy, it is very challenging to absolutely quantify iron content and cell number. Quantifying signal loss is not straightforward because the relationship between signal loss and iron concentration is not linear and because the presence of iron produces a localized disruption to the magnetic field homogeneity, resulting in a region of signal loss that extends far beyond the actual boundary of the cells. This is a susceptibility-related artifact known as the blooming effect. The size of the blooming artifact depends on several factors including magnetic field strength, type of iron particle, amount of iron particle uptake by the cell, the pulse sequence and pulse sequence parameters.^39^ Every SPIO positive DC within the node will contribute to the region of signal loss and, depending on the size of the signal voids, signal loss from multiple cells may overlap and cause complete signal loss in areas of accumulation. This is important when quantifying the total area of signal loss. The same number of cells, if dispersed in a larger region may result in a larger value for the signal void volume.^40^

In contrast, quantification of the MPI signal is directly related to the amount of iron in the ROI. The main advantage of MPI is the ability to directly quantify iron from the images. This can subsequently be used, along with a measurement of the amount of iron/cell, to estimate cell number. In this study MPI allowed for the estimation of the number of migrated DC in lymph nodes. To the best of our knowledge MPI has not yet been applied to imaging of DC. In our true migration experiment (Table 1), six mice were injected with either 300K (n = 3) or 500K (n = 3) Synomag-D+ DC. pLN signal was detected, and cell number quantified, *in vivo* in half of the target nodes. There was variability in the iron content quantified in nodes and the migrated cell numbers estimated from the MPI values. This is expected since there is biological variability in the mouse lymphatic system and in migration efficiency. MPI signal was not detected in the pLN of all mice. This may be explained by cell migration being below our detection threshold. Only 3-5% of injected DC migrate to a target lymph node post administration and this may be lower since varying numbers of DC are known to remain at the injection site, lose viability, and be cleared by infiltrating macrophages within 2 days.^41^ The inability to separate and resolve pLN signal from the larger footpad signal likely also contributes to our detection limits. Overall, this investigation provides the proof-of-principle concept that MPI can be used as a modality to track and quantify DC migration.

The main challenge for quantification of our MPI data was the isolation of neighboring strong and weak MPI signals, as shown in Figure 3. Detection of pLN signal was challenging *in vivo* where the larger footpad signal from remaining cells partially obscured the weaker signal from pLNs, preventing quantification in some mice. For this type of study, a wide dynamic range of signal is expected with high iron concentrations at the injection sites and low iron concentrations at migratory sites. These varying signal intensities contained in a small FOV result in the lower signal being shadowed by higher signal as shown in our mock migration experiment. GI signal was another challenge as a source of high signal, although we were able to mitigate GI signal by overnight fasting of animals with laxatives.

Several studies have reported on the issue of resolving a wide range of differing iron concentrations.^42–44^ Most recently, Boberg et al. discussed the idealized dynamic range of single sample and single iron concentration scenarios versus the effective dynamic range when there are multiple samples of different iron concentrations within the FOV.^17^ They found that the presence of high particle concentrations within the FOV can significantly reduce the dynamic range that the system is able to resolve. To show this reduced dynamic range, they used a realistic rat phantom containing kidneys as a low concentration source (1.5 mmol_Fe_L^−1^) and vessels as a high concentration source (100 mmol_Fe_L^−1^). Imaged separately, the kidneys and vessels were visible on their own. Imaged together, the kidneys were not visible next to the higher iron concentration of the vessels. As a solution, they proposed a two-step reconstruction algorithm to increase the dynamic range and reduce image artifacts. Herz et al discuss the issue as a function of the limited dynamic range of the analog to digital converter (ADC), with additional effects from reconstruction, discretization of the PSF, and the ADC not being used in the entire dynamic range.^44^ They introduce a selective signal suppression technique for traveling wave MPI that selects areas with low SPIO concentration to focus on in a large range of tracer concentrations. Alternatively, increasing the gradient strength could improve resolution of signals; however, this comes at the cost of sensitivity and longer imaging times.^45,46^ Changing the FOV to exclude the high signal source is another potential solution. Practically, this is challenging in cases where the exact location of signal is unknown, such as cell tracking where cells are dispersed, since image artefacts are caused by iron contained on or near the edge of the FOV, resulting in a large negative signal region.^47,48^

MPI does have some other limitations for tracking cells *in vivo*. Compared to iron-based MRI cell tracking, MPI has much lower resolution. This can make it challenging to pinpoint where the signal is located, especially without prior knowledge. It is expected that the ongoing development of MPI tailored SPIOs will substantially improve image resolution.^25,46,49–53^ To localize the MPI signal requires the acquisition of anatomical images and registration to MPI. Currently, our MPI system does not permit the acquisition of anatomical images though newer Momentum MPI systems are available with an integrated CT module. Finally, quantification of cell number can only be determined in experiments where SPIO is used to pre-label cells prior to their administration and the mass of iron per cell is known.

## Conclusion

Ideally, non-invasive imaging will be used as a biomarker for predicting the efficacy of DC immunotherapy. Quantification of the number of DC in lymph nodes by MPI could serve as an *in vivo* migration assay which could be used to determine whether more, or less, DC reach the target lymph nodes under different test conditions, such as: injection route, site of injection, timing of injection, maturation cocktails, cell number injected and antigen loading. Understanding the migration and accumulation of DC will provide valuable insight into the further development of cell-based cancer vaccines.

