## Supplementary Figure 1 for "Magnetic Particle Imaging is a sensitive *in vivo* imaging modality for the quantification of dendritic cell migration"

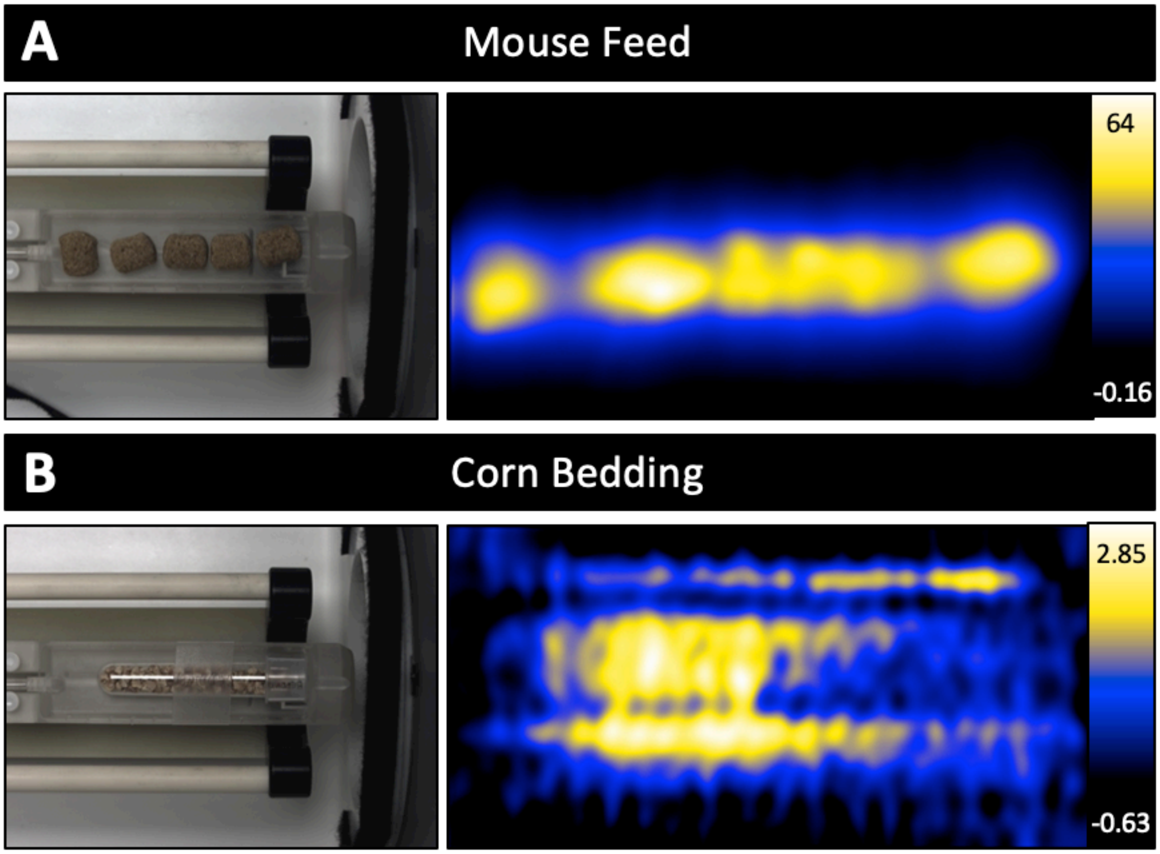


Supplementary Figure 1: MPI of 2D high sensitivity isotropic (3.0 T/m gradient) images of mouse feed (A) and corn cage bedding (B), both as sources of MPI signal contributing to GI signal in mice.
